# A highly contiguous, scaffold-level nuclear genome assembly for the Fever tree (*Cinchona pubescens* Vahl) as a novel resource for research in the Rubiaceae

**DOI:** 10.1101/2022.04.25.489452

**Authors:** Nataly Allasi Canales, Oscar A. Pérez-Escobar, Robyn F. Powell, Mats Töpel, Catherine Kidner, Mark Nesbitt, Carla Maldonado, Christopher J. Barnes, Nina Rønsted, Natalia A. S. Przelomska, Ilia J. Leitch, Alexandre Antonelli

## Abstract

**Background:** The Andean Fever tree (*Cinchona* L.; Rubiaceae) is the iconic source of bioactive quinine alkaloids, which have been vital to treating malaria for centuries. *C. pubescens* Vahl, in particular, has been an essential source of income for several countries within its native range in north-western South America. However, an absence of available genomic resources is essential for placing the *Cinchona* species within the tree of life and setting the foundation for exploring the evolution and biosynthesis of quinine alkaloids.

**Findings:** We address this gap by providing the first highly contiguous and annotated nuclear and organelle genome assemblies for *C. pubescens*. Using a combination of ∼120 Gb of long sequencing reads derived from the Oxford Nanopore PromethION platform and 142 Gb of short-read Illumina data. Our nuclear genome assembly comprises 603 scaffolds comprising a total length of 904 Mb, and the completeness represents ∼85% of the genome size (1.1 Gb/1C). This draft genome sequence was complemented by annotating 72,305 CDSs using a combination of *de novo* and reference-based transcriptome assemblies. Completeness analysis revealed that our assembly is moderately complete, displaying 83% of the BUSCO gene set and a small fraction of genes (4.6%) classified as fragmented. Additionally, we report *C. pubescens* plastome with a length of ∼157 Kb and a GC content of 37.74%. We demonstrate the utility of these novel genomic resources by placing *C. pubescens* in the Gentianales order using additional plastid and nuclear datasets.

**Conclusions:** Our study provides the first genomic resource for *C. pubescens*, thus opening new research avenues, including the provision of crucial genetic resources for analysis of alkaloid biosynthesis in the Fever tree.

## Data Description

### 1.1 Background

The Andes biodiversity hotspot hosts over 28,000 species [1], of which 3,805 benefit humanity [2]; unfortunately, nuclear genomic resources are only available for a limited number of such diversity (179 spp – Genomes NCBI database accessed on 26 May 2022). The fever tree (*Cinchona* L., Rubiaceae) is a genus comprising 24 species native to the Eastern slopes of the Andes mountain range in South America ([3,4]; Fig. 1) and one of the most economically important genera in the family, second only to coffee [5]. The genus is widely known as the source of at least 35 quinine alkaloids (quinolines), which alleviate the fever symptoms associated with malaria [6]. As such, fever trees have played a crucial role in the economies and livelihoods of people worldwide for centuries [7,8].

**Figure 1.** A. *Cinchona* trees in the Andean cloud forest. B. The *C. pubescens* specimen studied in this work (CP9014) growing in the Temperate House at the Royal Botanic Gardens, Kew, UK. C. Inflorescence of *C. pubescens*, CP9014. D. *Cinchona* barks from the Economic Botany Collection, Kew, UK. E. Distribution map of the *Cinchona* genus across the American continent shown in blue dots, modified from Maldonado et al. 2015 [75].

Despite this genus’s tremendous historical and economic importance, DNA sequence datasets for *Cinchona* are relatively meagre, limited to 252 DNA Sanger sequences available in the NCBI repository (accessed on May 17, 2021; [9]). More importantly, no nuclear and organellar reference genomes exist for any species of the genus. As such, important fundamental and applied questions – such as the mode and tempo of evolution of the fever tree or the genetic pathways responsible for quinine alkaloid production – remain elusive. Previous phylogenetic studies of the Rubiaceae family, specifically of the Cinchonoideae subfamily where the Cinchoneae tribe is, are based on just a handful of nuclear (ITS) and plastid (*mat*K, *rcb*L, *rps*16, *trn*L-F) data sets. They show an unresolved polytomy between the tribes and the seven genera of the Cinchoneae tribe that have so far been included in more specific studies [10,11] (including the genus *Cinchona*, which shows very unclear relationships). Furthermore, studies that examine the relationships between species of this genus are equally scarce [7,8]. A recent genome-wide phylogenetic tree for the order Gentianales [12] provided strong support for *C. pubescens* as a sister to *Isertia hypoleuca*, but the sampling was exclusively at the genus level and therefore did not include any other species of *Cinchona* nor other genera in tribe Cinchoneae.

The production of alkaloids is highest in *C. calisaya*, also known as yellow bark [13][14]. However, several species in the genus have historically been harvested to provide sources of quinine alkaloids, one of the most traded natural products, resulting in significant reductions in their natural ranges and population size [15,16]. Among them, *C. pubescens* or red Cinchona bark is now widely cultivated throughout the tropics, with some instances where the species has escaped cultivation and become invasive [17]. Extensive research on the structure, abundance, and chemical composition of quinine alkaloids in *Cinchona* has been conducted [18], revealing the further potential for novel drug discovery. However, the identity of the genes involved in the synthetic pathway of quinine alkaloids remains elusive.

Nuclear genome assemblies are critical to our understanding of the origin and domestication of useful plants and are a cornerstone resource for breeders [19–21]. Here, we present the first high-quality draft nuclear and plastid genomes of *C. pubescens*, which is characterised by having a genome size of 1.1 Gb (1C, this study) and a chromosome number of 2*n*=34. The assemblies were generated using a combination of extensive long-read Nanopore (∼218x) and short-read Illumina paired-end read datasets (∼300x) jointly with state-of-the-art genome assemblers, resulting in a reference genome for which contiguity and quality are comparable to, or even higher [22] than in the three previously published genome assemblies in Rubiaceae, namely for *Chiococca alba [22], Coffea canephora* [23], and *Coffea arabica [24]*. The plastid genome from short-reads of *C. pubescens* had a length of 156,985 bp and a GC content of 37.74%, very similar to other Rubiaceae plastid genomes [22,25]. Lastly, we demonstrate the utility and reliability of our resources by constructing nuclear and plastid phylogenomic frameworks of *C. pubescens*.

### 1.2 Sampling and genomic DNA and RNA sequencing

We sampled leaves from a single *Cinchona pubescens* individual propagated vegetatively from a tree collected in Tanzania in 1977 and cultivated in the Temperate House of the Royal Botanic Gardens, Kew (RBG Kew), UK (Accession Number 1977-69; a voucher was also prepared which is deposited in the RBG Kew herbarium [**K**]). DNA was extracted from fresh tissue using two different protocols to produce paired-end Illumina and native Nanopore libraries. For Illumina DNA library preparations, we used 1000 mg of starting material that was first frozen with liquid nitrogen and subsequently ground in a mortar. The Qiagen DNeasy (Qiagen, Denmark) plant kit was used to extract DNA from the ground tissue, following the manufacturer’s protocol. We built the libraries using the Illumina TruSeq PCR-free library (NEX, Ipswich, MA, USA) following the manufacturer’s protocol, by first assessing the DNA quantity and quality using a Nanodrop fluorometer (Thermo Scientific, Denmark) and then fragmenting oligonucleotide strands through ultrasonic oscillation using a Covaris ME220 (Massachusetts, USA) device to yield fragments with an average length of 350 bp. Then we sequenced the paired-end 150 bp libraries using the HiSeq X Ten chemistry. For transcriptome library preparations, total RNA was extracted from 1000 mg of frozen-ground leaf, young bract, mature bract, flower anthesis, flower bud (older), flower bud (young), leaf bud and young leaf tissue using the TRIZoL reagent (Thermo Fisher Scientific, Denmark) following the manufacturer’s protocol. Illumina library preparation and sequencing were conducted by Genewiz GmbH (Leipzig, Germany).

The Nanopore sequencing data were generated and base called as part of Oxford Nanopore’s London Calling 2019 conference [26]. For Nanopore library preparation, 1000 mg of leaf tissue was frozen and ground with a mortar and pestle. The lysis was carried with Carlson lysis buffer (100 mM Tris-HCl, pH 9.5, 2% CTAB, 1.4 M NaCl, 1% PEG 8000, 20 mM EDTA) supplemented with β-mercaptoethanol. The sample was extracted with chloroform and precipitated with isopropanol. Finally, it was purified using the QIAGEN Blood and Cell Culture DNA Maxi Kit (Qiagen, UK). Size selection was performed using the Circulomics Short Read Eliminator kit (Circulomics, MD, USA) to deplete fragments below 10 kb. DNA libraries were prepared using the ONT Ligation Sequencing Kit (SQK-LSK109, Oxford Nanopore Technologies, UK). During sequencing on the PromethION platform, re-loads were performed when required. Though yield was slightly lower in sequencing for these re-loaded samples (over 50 Gb in 24 hours), the read N50 was over 48 kb (up from 28 kb without size selection).

### 1.3 Estimation of genome size

To accurately determine the genome size of *C. pubescens*, we followed the one-step flow cytometry procedure [27], with modifications as described in Pellicer et al. [28]. Freshly collected tissue from the same individual sampled for DNA and RNA sequencing was measured together with *Oryza sativa* L. ‘IR-36’ as the calibration standard using general-purpose buffer” (GPB) [29] supplemented with 3% PVP-40 and β-mercaptoethanol [28]. The samples were analysed on a Partec Cyflow SL3 flow cytometer (Partec GmbH, Münster, Germany) fitted with a 100 mW green solid-state laser (532 nm, Cobolt Samba, Solna, Sweden). Three replicates were prepared and the output histograms were analysed using the FlowMax software v.2.4 (Partec GmbH, Münster, Germany). The 1C-value of *C. pubescens* was calculated as (Mean peak position of *C. pubescens*/Mean peak position of *O. sativa*) × 0.49 Gb (=1C value of *O. sativa*) [30] and resulted in a 1C-value of 1.1 Gb. Additionally, using the full Illumina short-read dataset, we additionally implemented a k-mer counting method to characterise the genome in Jellyfish v.2.2.10 [31] setting a kmer size of 21. We used GenomeScope to visualise the kmer plot [32]. However, we did not deem the kmer counting method sufficiently accurate for genome size estimation but reported the estimated genome-wide heterozygosity rates output by GenomeScope as 0.869-0.889%.

### 1.4 Short read data processing of the chloroplast genome assembly

Sequencing of the DNA Illumina library generated 428M paired-end reads, representing 128.4 Gb of raw data. RNA sequencing produced 385M paired-end reads, representing 115.5 Gb (Table 1). The quality of the raw reads was assessed using the FastQC software [33], and quality trimming was conducted using the software AdapterRemoval2 v.2.3.1 [34]. Here, bases with Phred score quality <30 and read lengths <50 bp were removed together with adapter sequences. The final short-read dataset was 131 Gb and contained 384,626,011 paired reads, which corresponds to an estimated 464.8x coverage (based on the genome size of 1.1 Gb/1C, see *Estimation of genome size*).

**Table 1.**
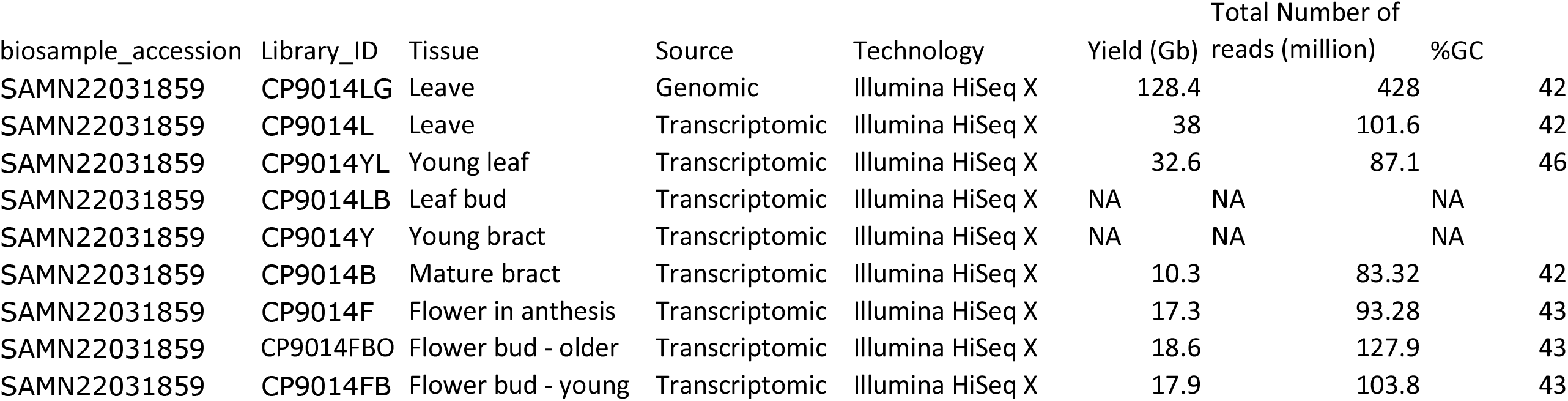
Summary table for the Illumina WGS and RNA-Seq libraries.

**Table 2.** Summary assembly statistics for *C. pubescens* using SMARTdenovo and RACON.

|  | SMARTdenovo | RACON |
| --- | --- | --- |
| Size_includeN | 903037179 | 9.04E+08 |
| Size_withoutN | 903037179 | 9.04E+08 |
| Seq_Num | 603 | 603 |
| Mean_Size | 1497574 | 1499914 |
| Median_Size | 801066 | 802662 |
| Longest_Seq | 14628764 | 14747124 |
| Shortest_Seq | 33171 | 25882 |
| GC_Content (%) | 33.17 | 33.07 |
| N50 | 2783363 | 2802128 |
| L50 | 93 | 92 |
| N90 | 682446 | 684435 |
| Gap (%) | 0 | 0 |

**Table 3.** Annotation metrics summary statistics for the nuclear annotation of *C. pubescens*.

|  |  |
| --- | --- |
| Number of gene models | 72305 |
| Minimum gene length | 275 |
| Maximum gene length | 65857 |
| Average gene length | 4579.5 |
| Number of exons | 339698 |
| Average number of exons per gene model | 4.7 |
| Average exon length | 298.3 |
| Number of transcripts | 72305 |
| Average number of transcripts per gene model | 1 |
| Number of gene models less than 200bp length | 0 |

The plastid genome of *C. pubescens* was assembled using only short reads, as there were some discrepancies using the hybrid dataset. The toolkit GetOrganelle v.1.7.5, was used with the parameters suggested for assembling plastid genomes in Embryophyta (i.e., parameters *-R* 15, *-k* 21,45,65,85,105, *-F* embplant_pt). GetOrganelle produced a single linear representation of the *C. pubescens* plastid genome, with a length of 156,985 bp (Fig. 2) and a GC content of 37.74%. These values are very similar to those reported for the *Coffea arabica* plastid genome, which is reported to be 155,189 bp in length and has a GC content of 37.4% [25].

**Figure 2.**
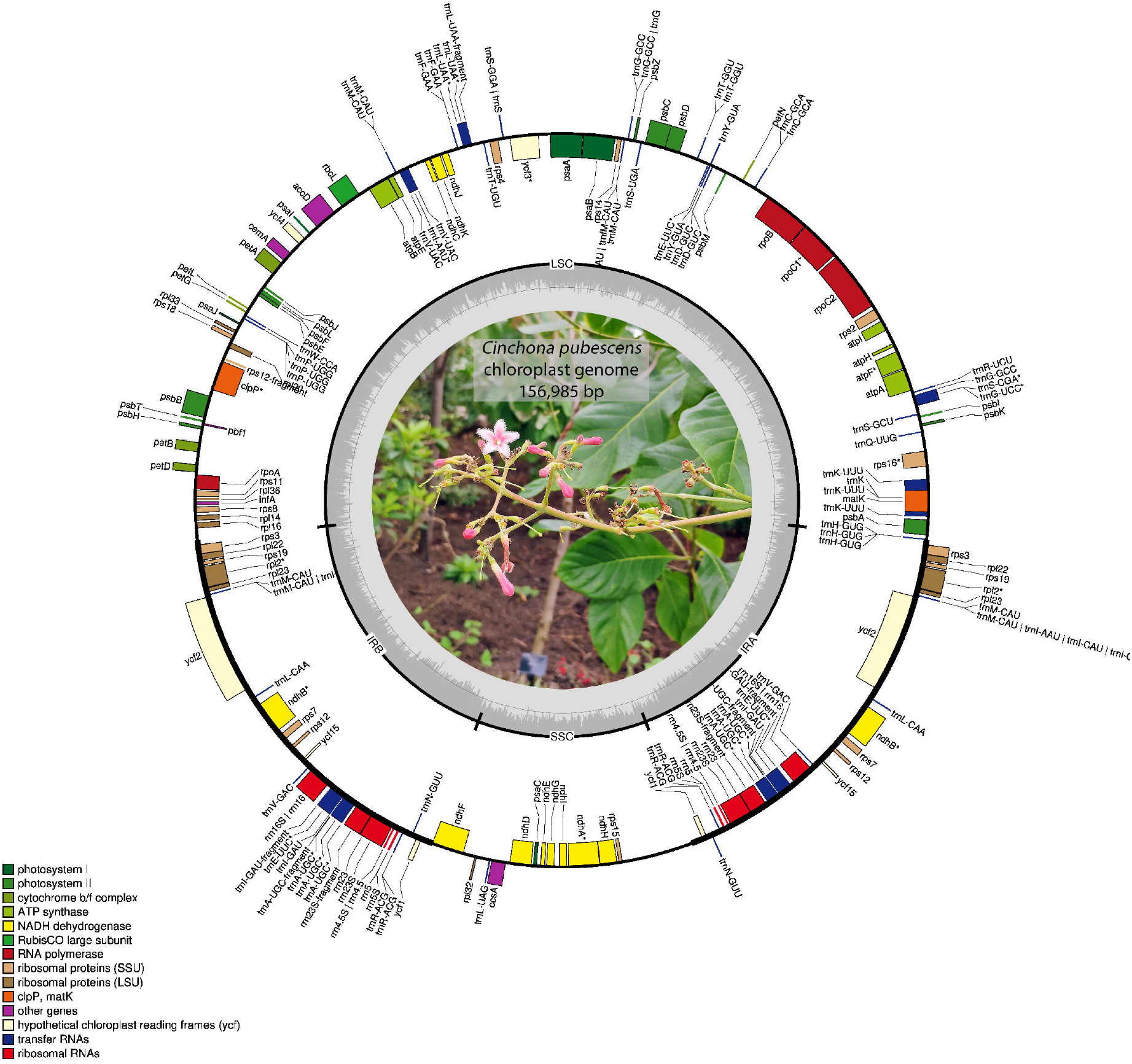
Annotated *C. pubescens* plastome. Genes displayed on the inside of the circle are transcribed clockwise, while genes positioned on the outside are transcribed counter clockwise.

We annotated the plastid genome assembly of *C. pubescens* in CHLOROBOX [35], which implements GeSeq [36], tRNAscan-SE v2.0.5 [37], and ARAGORN v1.2.38 [38]. CHLOROBOX annotations indicated that the *C. pubescens* genome has the typical angiosperm quadripartite structure, i.e., Inverted Repeat (IRa and IRb) (each 27,502 bp long), the Small Single Copy (SSC) region (18,051 bp), and the Large Single Copy (LSC) region (83,930 bp). We predicted 128 genes, of which 34-37 were tRNA (tRNAScan-SE and ARAGORN, respectively), 81 CDSs, and four ribosomal RNAs (rRNAs). The junction between SSC-IRa and LSC-IRa contains the *ycf1* pseudogene and *rps3* gene, respectively. Similarly, the junction between IRb-SSC and LSC-IRb contains the *ycf1* pseudogene and *rsp3* gene, respectively (Supp. Fig. 1). The final structural features of the *C. pubescens* plastid genome were generated using OGDRAW v. 1.3.1 [39] (Fig. 2) and edited manually. Finally, the quality of the plastid genome assembly was estimated by mapping the Illumina DNA short reads to the newly assembled genome using the bam pipeline in Paleomix [40], where we used BWA [41] for alignment, specifying the backtrack algorithm, and filtering minimum quality equal to zero to maximise recovery. After PCR duplicate filtering, the coverage of unique hits was 7,960x.

### 1.5 Long-read nuclear genome assembly, quality assessment and ploidy levels

The quality and quantity of the PromethION sequencing output conducted across four flow cells were evaluated in NanoPlot v.1.82 [42] independently for each flow cell, using as input the sequencing summary report produced by Guppy v3.0.3. Overall, the average read length, Phred score quality and N50 following base calling with Guppy v3.0.3, and the High Accuracy model reached values of ∼19,000 bp, 9, and ∼46,000 bp, for mean read length, mean read quality and read length N50, respectively (Supp. Tab. 1). A total of 13,252,640 quality-passed reads were produced, representing ∼262 Gb and providing a theoretical genome coverage of ∼218x. To assemble the raw Nanopore reads into scaffolds, we first corrected and trimmed the quality-passed reads using the software CANU v.1.9 [43] in correction and trimming mode with the following parameters: *genomeSize* = 1.1g, *-nanopore-raw*. This step generated a total of 1,265,511 reads, representing c. 89 Gb, or a theoretical genome coverage of 74x. Next, the corrected/trimmed reads were used as input into SMARTdenovo v.1.0 [44], using the following parameters: *-c* 1 (generate consensus mode), -*k* 16 (k-mer length) and *-J* 5000 (minimum read length). This step produced an assembly composed of 603 scaffolds with an N50 of 2,783,363 bp, representing ∼904 Mb (∼82% of the genome size; Tab. 1). Lastly, a round of scaffold correction was implemented in RACON v.1.4.3 [45] using as input the corrected Nanopore reads generated by CANU and an alignment SAM file produced by mapping the trimmed DNA Illumina reads against the assembly produced by SMARTdenovo. The alignment file was produced by Minimap2 v.2.18 [46] using the “accurate genomic read mapping” settings designed to map short-read Illumina data (flag *-ax*). RACON was executed using an error threshold of 0.3 (*-e* flag), a quality threshold of 10 (*-q*), and a window length of 500 (*-w*). The corrected assembly differed little compared with the raw assembly produced by SMARTdenovo (Tab. 1).

We followed a two-pronged approach to assess the quality of our corrected nuclear genome assembly by i) evaluating the proportion of Illumina reads that mapped against our new genome assembly using as input the alignment file (SAM) generated by Minimap2 and computing coverage and mean depth values per scaffold, as implemented in the function *view* (flag –F 260) of the software Samtools v1.12 [47]; and ii) estimating the completeness of the genome as implemented in the software BUSCO v.5.2 and using the viridiplantae_odb10 [48]. A total of 827,098,761 reads were mapped against the corrected genome assembly, representing 99% of the trimmed reads used as input (241,498,983). Mean coverage and read depth ranged from 26-48x. The genome completeness analysis recovered a total of 92.4% conserved eudicot genes, of which 87.5% were single copy, 4.9% duplicated, and 5.6% fragmented. The remaining BUSCO genes were labelled as missing (2%). Taken together, our results suggest that our nuclear genome assembly presents high contiguity and quality with high completeness.

Lastly, to evaluate the ploidy levels of *C. pubescens* through the newly assembled genome, we computed allele frequencies from reads mapped against two scaffolds, “utg 230” and “utg2” derived from our genome assembly, covering 9,568,509 bp (∼106x coverage) and 14,628,764 bp (∼103x coverage), respectively. The reads were obtained from the mapping procedure conducted to assess the quality of the corrected nuclear genome assembly (see above). We relied on the software ploidyNGS to compute allele frequencies [49], using the –g option (i.e. guess ploidy levels), a maximum read depth of 100 option (-d 100) and a maximum allele frequency of 0.95 (-m 0.95). Our analysis revealed that the genome of *C. pubescens* is diploid (Fig. 3) as inferred by the comparison of Kolmogorov-Smirnoff distances between the allele frequencies computed from our read mappings and those derived from simulated data [49].

**Figure 3.**
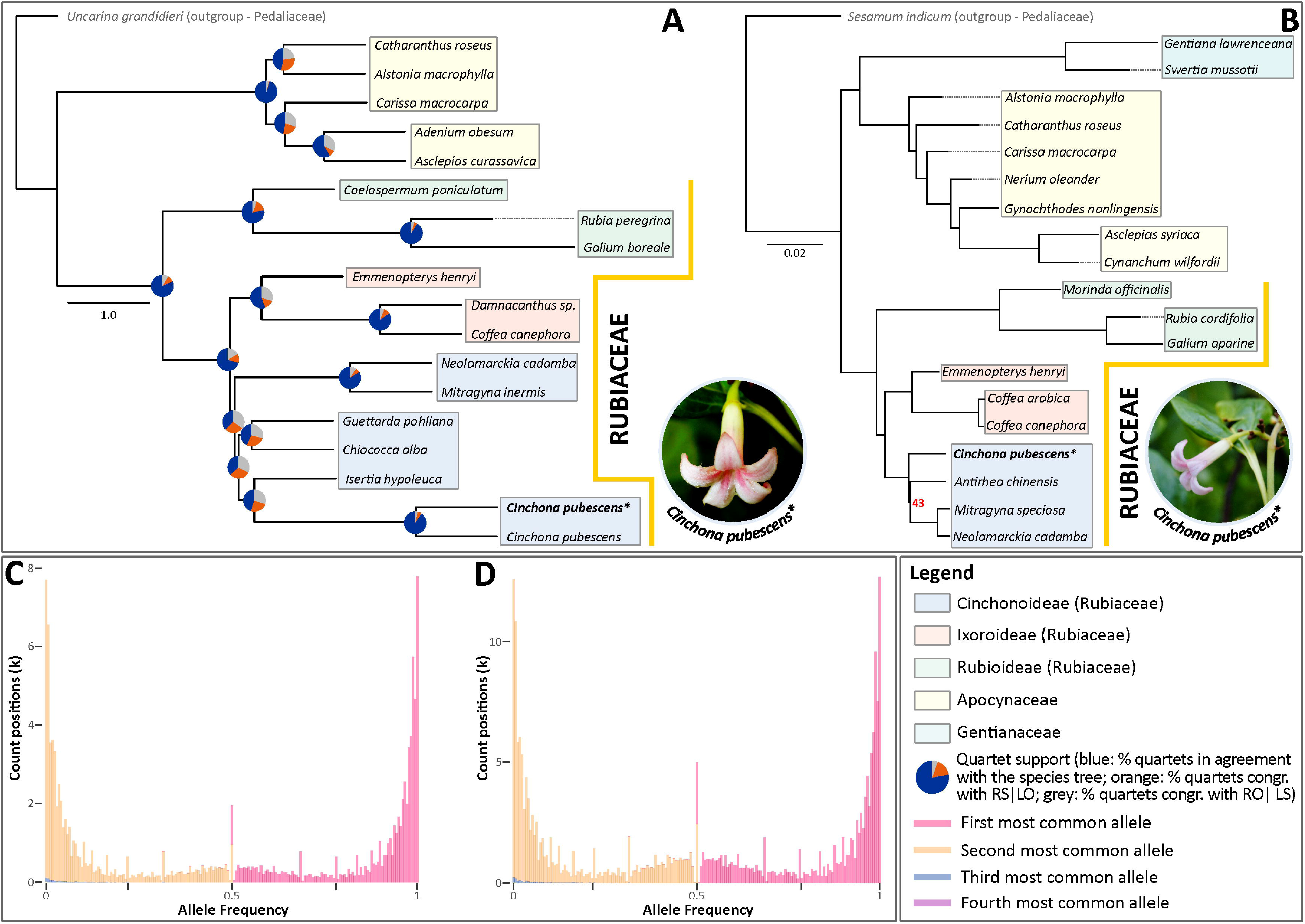
**A**. The coalescent-based species tree estimation of the Gentianales order is inferred from low copy nuclear gene trees. Pie charts positioned on the nodes represent the percentage of the gene tree quartets that agree with the topology of the main species tree (blue) and the other two alternative gene tree quartets (orange: second child [R], sister group [S] | first child [L], any other branch [O]; and grey: RO|LS). The genomic data of *Cinchona pubescens* terminal marked with “*” was newly produced in this study. **B**. Phylogenetic tree showing the relationships of twenty Gentianales species built from the whole plastid genome. Unless shown, numbers on the branches represent Likelihood Bootstrap Percentages of 100. The coloured boxes indicate the subfamily/family that sampled terminals belong to. Allele frequencies for the fourth most common alleles, derived from reads mapped against the scaffolds “utg230” (C) and “utg2” (D). Note that frequency distributions of the first and second most frequent alleles are either skewed towards 0 or 1, denoting homozygote variants, and 0.5, denoting heterozygote, diploid variants. (Inset): Frontal (left) and lateral view (right) of the sequenced specimen (ID 1977-69) of *Cinchona pubescens* (photos: O. A. Pérez-Escobar).

### 1.6 Transcriptome assembly, candidate gene annotation, and quality assessment

To produce a comprehensive database of assembled transcripts, we generated reference-based and *de-novo* assemblies with the Trinity toolkit v. 2.8 [50] using the trimmed RNA-seq data. The reference-based assembly was conducted using as input the aligned RNA-seq trimmed reads against our new reference genome as produced by aligner STAR v.2.9 [51] with default settings, and a maximum intron length of 57,000 as estimated for *Arabidopsis thaliana* (flag *--genome_guided_max_intron*). The *de-novo* transcriptome assemblies were also produced using the default settings of Trinity and the trimmed RNA-seq reads as input. A comprehensive database of *de-novo* and reference-based assembled transcriptomes was compiled with the software PASA v.2.0.2 [52], using the following parameters: --*min_per_ID* 95, and --*min_per_aligned* 30.

To assess the completeness of the *de-novo* transcriptome assembly, we used BUSCO v.5.12 and the representative plant set viridiplantae_odb10, which currently includes 72 species, of which 56 are angiosperms. Our assembled transcriptome captured 92.7% (394/425) of the BUSCO set as complete genes, of the remainder, 3.1% of the genes were fragmented, and 4.2% were missing.

We predicted the structure and identity of the genes in the nuclear genome using the comprehensive transcriptome assembly compiled with PASA. For this purpose, we used AUGUSTUS v3.3.3 [53] for a combination approach of *ab initio* and transcript evidence-based on RNA-seq data. As AUGUSTUS considered the transcripts’ evidence as Expressed Sequence Tag (EST), we first generated hints from the transcriptome data by aligning the transcripts to the genome using BLAT v 3.5 [54]. Then, we set the hint parameters to rely on the hints and anchor the gene structure. We predicted 72,305 CDSs using the hints and tomato (*Solanum lycopersicum* L.) as the reference species. The completeness of the CDSs was estimated using BUSCO v.5.12 and vidiriplantae odb10 [48] as reference: 68.4% represented complete BUSCOs, 63.5% were single-copy, 26% were fragmented and 5.6% were missing. We summarized the metrics of the CDSs statistics with GenomeQC (Table 1) [55]

### 1.7 Nuclear and plastid phylogenomics of *Cinchona*

We verified the nuclear genome’s phylogenetic placement using the reference sequences of the 353 low-copy nuclear genes that are conserved across angiosperms from the Plant and Fungal Trees of Life project [56]. Here, we sampled the gene sequences for the 18 taxa from the Gentianales, which included another *C. pubescens* from that study (Supp. Tab. 2) that are publicly available in the Tree of Life Explorer [57] hosted by the Royal Botanic Gardens, Kew. To include the *C. pubescens* of this study in the analysis of the 353 low copy nuclear genes of selected Gentianales, we then retrieved these genes from our RNA-seq data using the pipeline HybPiper v.1.3.1 [58]. Given the abundance of RNA-seq read data, to render the gene retrieval tractable, as input for HybPiper we used a subsample of the trimmed read data, as implemented in the software seqtk [59]. The gene sequences produced by HybPiper were aligned with the data for 19 selected Gentianales species using MAFFT v7.453 [60] and then they were concatenated into a supermatrix for phylogenomic analyses.

We implemented the maximum likelihood approach using RAxML-HPC V.8 [61] with a GTRGAMMA substitution model for each gene and a rapid bootstrap analysis with 500 replications. Then we filtered the bipartition trees that had >=20% support using Newick utilities [62]. The resulting trees were rooted using phyx v1.2.1 [63], setting *Uncarina grandidi*eri (Baill.) Stapf (Lamiales) as the root. To estimate the species tree from the gene trees, we used the coalescent approach with ASTRAL 5.6.1 to calculate the quartet scores, which is the number of quartet trees present in the gene trees that are also present in the species tree. Q1 shows the support of the gene trees for the main topology, q2 shows the support for the first alternative topology, and q3 shows the support for the second alternative topology [64]. We incorporated these scores into the species tree with an R script [65]. All trees were visualized with FigTree v.1.4.4 [66].

In the nuclear phylogenomic tree resulting from the 353 low copy nuclear genes (Fig. 3), *C. pubescens* clusters within the Cinchonoideae, which is more closely related with the Ixoroideae group than with Rubioideae. Most nodes are highly supported by quartet scores, showing that a large proportion of the gene trees agreed with the species tree.

For the plastid phylogeny, we used *Sesamum indicum* L. as an outgroup from the Lamiids cluster [67]. We performed maximum likelihood using the complete plastid genomes of the 20 species available to date in the Gentianales. All the plastid genomes we analysed had the classic quadripartite genomic structure, although some Rubiaceae species show the tripartite structure [68]. We aligned the 20 Gentianales (Supp. Tab. 3) plastid genomes with MAFFT v7.427 using the default parameter settings to perform the multiple sequence alignments. Then we estimated the phylogenetic tree with the maximum likelihood (ML) approach using the GTRCAT model RAxML-HPC v.8. We conducted heuristic searches with 1000 bootstrap replicates (rapid bootstrapping and search for the best-scoring ML tree). Both analyses were performed on the Cipres Science Gateway [69].

As with the nuclear tree, the plastid trees were also clustered at the subfamily level, recovering the Cinchonoideae, Ixoroideae, and Rubioideae as natural groups, alongside the two species belonging to Pedilaceae used as outgroups for the phylogenetic analysis. For the plastid data, the vast majority of nodes were strongly supported (16 had 100% support and all but one node had 100% support). However, we found *Gynochthodes nanlingensis* (Y.Z.Ruan) Razafim. & # x 0 0 2 6; B.Bremer (Rubioideae) to cluster with other Apocynaceae species. While the same result has previously been reported in other studies [70,71], it seems to be due to an erroneous DNA sequence attributed to *G. nanlingensis* or a misidentification of the voucher, so it is recommended this is thoroughly checked. Additionally, the ingroup showed that the Cinchonoideae and Ixoroideae subfamilies are sisters and form a clade while Rubioideae is placed as sister to this clade.

The placement of *C. pubescens* in the Cinchonoideae subfamily cluster using both the plastid and nuclear data presented in this study is consistent with previous taxonomic and phylogenetic studies [72] and gives support to the robustness of the assembled nuclear and plastid genomes. As potential future work, the Nanopore sequencing data could be re-base called using the latest algorithms from Oxford Nanopore to take advantage of recent developments in this area over the last few years which has seen continuous improvement in raw-read accuracy [73,74].

## 2. Conclusion

Using a combination of extensive short and long-read DNA datasets, we deliver the first highly contiguous and robust nuclear and plastid genome assemblies for one of the historically most traded and economically important *Cinchona* species, *C. pubescens*. The abundant genomic resources provided here open up new research avenues to disentangle the evolutionary history of the Fever tree.

In the short term, these genomic tools will significantly help to identify the genes involved in the biosynthetic pathways of quinine alkaloids synthesis, identify the underpinning genetic diversity of these genes both between and within species, and open doors on how the expression of these genes is regulated. Our nuclear scaffold-level and plastid genome assembly will enable future reference-guided assemblies, variant calling, and gene annotation to enhance functional analysis within the *Cinchona* genus, with the potential to further explore the quinine alkaloid biosynthetic pathway in-depth and hence enhance its potential for finding new medicinal leads to treat malaria.

## Data availability

The genome sequence data, and nuclear and plastid assemblies are available at the NBCI repository, under the BioProject number PRJNA768351.

## Author contributions

IJL, OAPE, NR, MT and AA conceived the study. MN and OAPE collected plant tissue. OAPE, IJL, RFP, MT and AA generated datasets. OAPE, NC, CK and MT conducted in-silico analyses. NC and OAPE wrote the manuscript, with contributions from all co-authors.

## Acknowledgements

We thank Jonathan Pugh, Vania Costa, and Simon Mayes for support and assistance during Nanopore sequencing preparation and Claes Persson for taxonomic advice. Two anonymous reviewers and the associate editor provided constructive feedback to this manuscript.

## Additional files

Supplementary Table 1. Summary statistics of the Nanopore reads.

Supplementary Table 2. Overview of the samples from the Tree of Life Explorer (Royal Botanic Gardens, Kew) that were used in the phylogenetic analysis to construct the coalescent tree.

Supplementary Table 3. Sample overview of the specimens and their accession numbers used to infer the phylogenetic tree built using plastid data.

Supplementary figure 1. Plastid genome visualization of junctions IRb-SSC (*ycf1*) and LSC-IRb (*rsp3*).

