## Supplementary figures and images for "A highly contiguous, scaffold-level nuclear genome assembly for the Fever tree (*Cinchona pubescens* Vahl) as a novel resource for research in the Rubiaceae"

### Fig1_Supp

*C. pubescens* plastid genome assembly

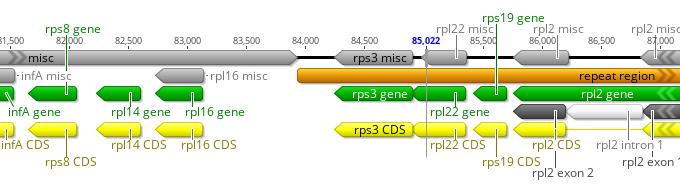

LSC-IRa

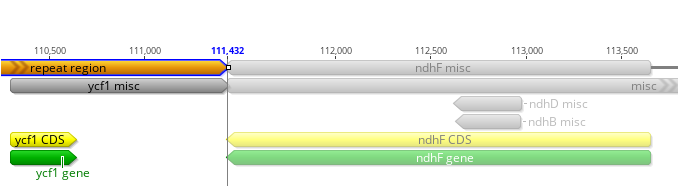

IRa-SSC

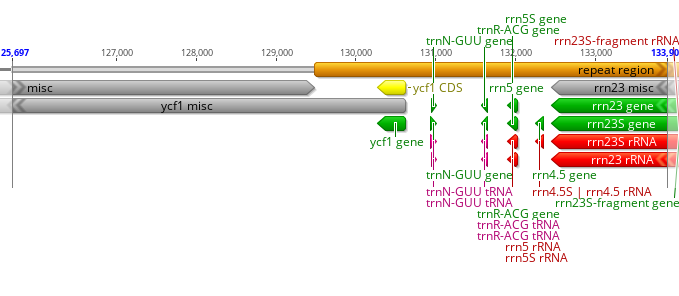

SSC- IRb

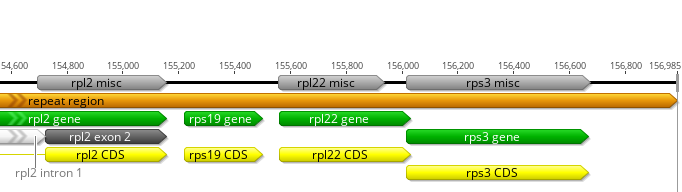

IRb-LSC

### Fig. 1

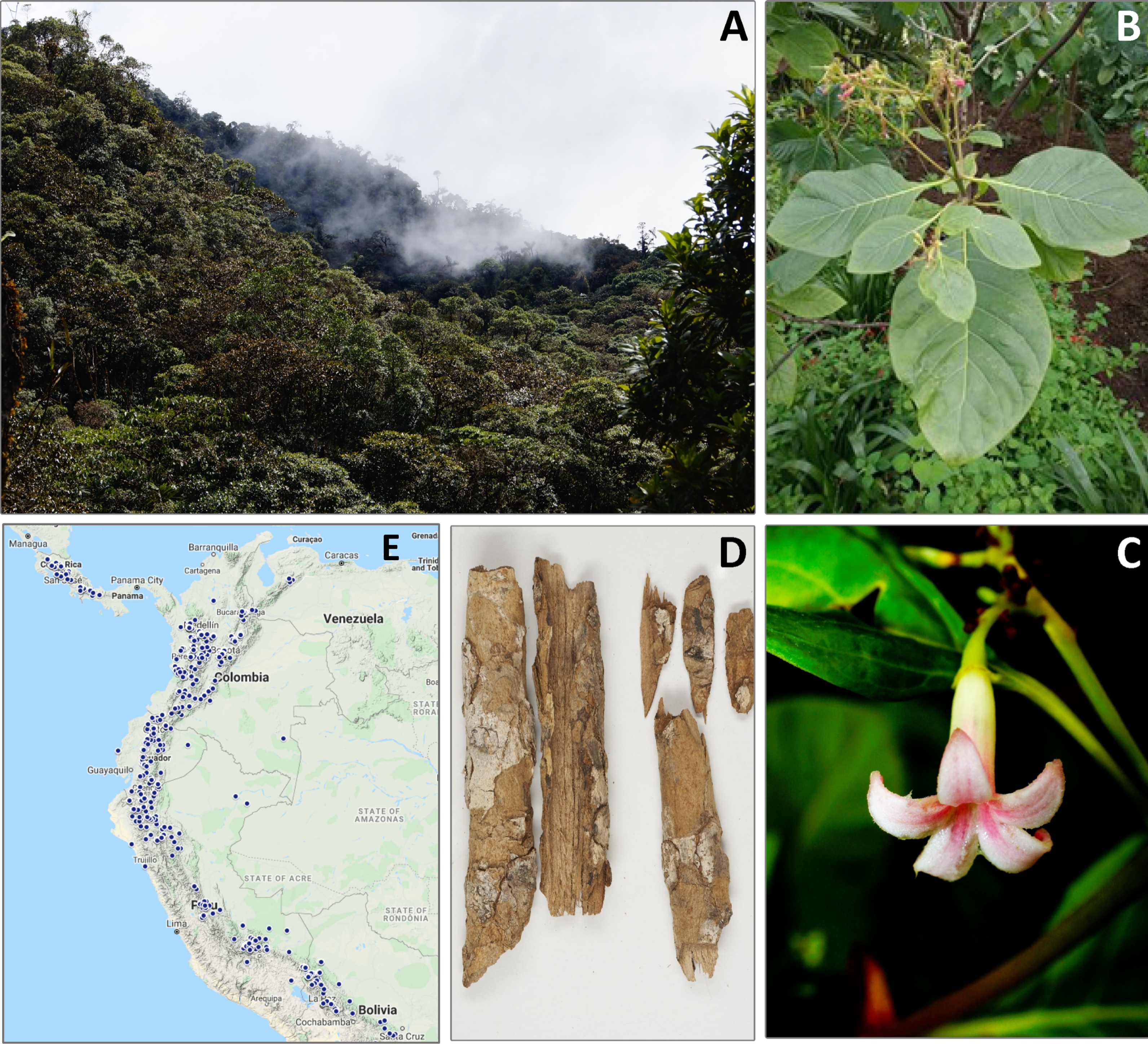
